# Exploiting KRAS-driven Ferroaddiction in Cancer Through Ferrous Iron-Activatable Drug Conjugates (FeADC)

**DOI:** 10.1101/2020.05.12.088971

**Authors:** Honglin Jiang, Ryan K. Muir, Ryan L. Gonciarz, Adam B. Olshen, Iwei Yeh, Byron C. Hann, Ning Zhao, Yung-hua Wang, Spencer C. Behr, James E. Korkola, Michael J. Evans, Eric A. Collisson, Adam R. Renslo

## Abstract

*KRAS* mutations cause a quarter of cancer mortality and most are undruggable. Several inhibitors of the MAPK pathway are FDA approved but poorly tolerated at dosages required to adequately extinguish RAS/RAF/MAPK signaling. We found that oncogenic KRAS signaling induces ferrous iron (Fe^2+^) accumulation early in and throughout KRAS-mediated transformation. We used an FDA-approved MEK inhibitor to produce a prototypical Ferrous Iron–Activatable Drug Conjugate (FeADC) which achieved potent MAPK blockade in tumor cells while sparing normal tissues. This innovation allowed sustainable, effective treatment of tumor bearing animals, with tumor-selective drug activation producing superior systemic tolerability. Ferrous iron accumulation is an exploitable feature of KRAS transformation and FeADCs hold promise for improving treatment of KRAS-driven solid tumors.

## Introduction

The uptake, transport, and storage of iron is tightly controlled in biology but often dysregulated in cancer. Transcriptional profiling has suggested a link between elevated iron levels and poor prognosis in in gliomas (Han et al., 2018), breast (Torti and Torti, 2013a) and prostate cancers (Tesfay et al., 2015), but clinical studies based on iron chelation have proved disappointing (Buss et al., 2003). New insights into iron homeostasis have been revealed through genome-wide association studies (GWAS) and the recent development of iron probes with oxidation-state specificity. A recent functional screen found that the pH-dependent unloading of Ferric iron from transferrin, its subsequent reduction and delivery of bioavailable ferrous iron (Fe^2+^) to the cell was the single indispensable function of the lysosome in cell proliferation (Weber et al., 2020). This remarkable finding was perhaps anticipated from studies of the bacterium *P. aeruginosa*, where genetic deletion of a bacterioferrireductase (limiting cytosolic Fe^2+^) produced acute iron starvation despite more than adequate Fe^3+^ bacterioferritin stores (Eshelman et al., 2017). Elevation of cytosolic Fe^2+^ may enable the survival of drug tolerant ‘persister’ cancer cells (Hangauer et al., 2017), which become uniquely reliant on the activity of GPX4, a lipid hydroperoxidase that protects from ferroptosis (Dixon and Stockwell, 2014; Stockwell et al., 2017). Taken together, these studies suggest that access to bioavailable Fe^2+^ (as opposed to total iron) is a key to cancer cell survival. The mechanisms by which oncogene-mediated transformation lead to such ‘ferroaddiction’ remain unknown.

Pancreatic ductal adenocarcinoma (PDA) is among the most aggressive and lethal solid tumors with a 5-year survival rate of ~9% (Siegel et al., 2018). Systemic therapies are only marginally effective and no targeted therapies against somatic aberrations exist in this disease. *KRAS* mutation is a cardinal feature of PDA. Constitutive RAS activation drives uncontrolled proliferation and enhances survival of cancer cells, but also sensitizes tumor cells to ferroptosis (Dixon et al., 2012). Here we explore the elevation of intracellular Fe^2+^ in mutant *KRAS*-driven tumor cells *in vitro* and *in vivo*, and target these tumors with a prototypical Ferrous Iron–Activatable Drug Conjugate (FeADC) that ablates MAPK signaling in tumor while sparing tissues that are dependent on physiologic RAS/MAPK signaling for tissue homeostasis, allowing a dramatically improved therapeutic window.

## Results

### A Ferrous Iron Signature is Prognostic in Pancreatic Adenocarcinoma

We first evaluated mRNA transcriptional data from 23 tumor types profiled by TCGA with a focus on Iron homeostatic programs using biological pathway activity (BPA) method (Ding et al., 2019) to explore the role of iron metabolism in cancer in an unbiased manner. The cancer types with the highest Z-scores for iron metabolism were all RAS-MAPK pathway-driven malignancies including: Glioblastoma Multiforme (GBM), Lung Adenocarcinoma (LUAD) and pancreatic ductal adenocarcinoma (PAAD) (**Figure 1A).***STEAP3* (a lysosomal ferrireductase) and *TFRC* (transferrin receptor) encode key genes in iron homeostasis. We compared *STEAP3* and *TFRC* mRNA levels to overall patient survival (OS) in PAAD, a disease with the highest rates of *KRAS* mutations. Both *STEAP3* and *TFRC* expression were associated with short OS by Cox regression (**Figure 1B**). Liver metastasis is a uniformly poor prognostic sign in PDA. We examined *STEAP3* expression in an independent primary and metastatic PDA dataset (Aguirre et al., 2018) and found *STEAP3* expression to be higher in liver metastases than in all other metastatic sites (p=0.0001, **Figure 1C**). Gallium-68 citrate is a clinically studied radiotracer that mimics Fe^3+^-loaded on transferrin and can quantify iron demand in cancer patients (Behr et al., 2018; Larson et al., 1980). To assess iron avidity in PDA, we conducted an exploratory human imaging study with Ga-68 citrate positron emission tomography (PET) in our PDA patients. Quantitative analysis of the imaging data showed that visceral and bony PDA metastases were highly avid for Gallium-68 compared to normal tissues, consistent with a Ferroaddiction phenotype **Figure 1D.**

**Figure 1.**
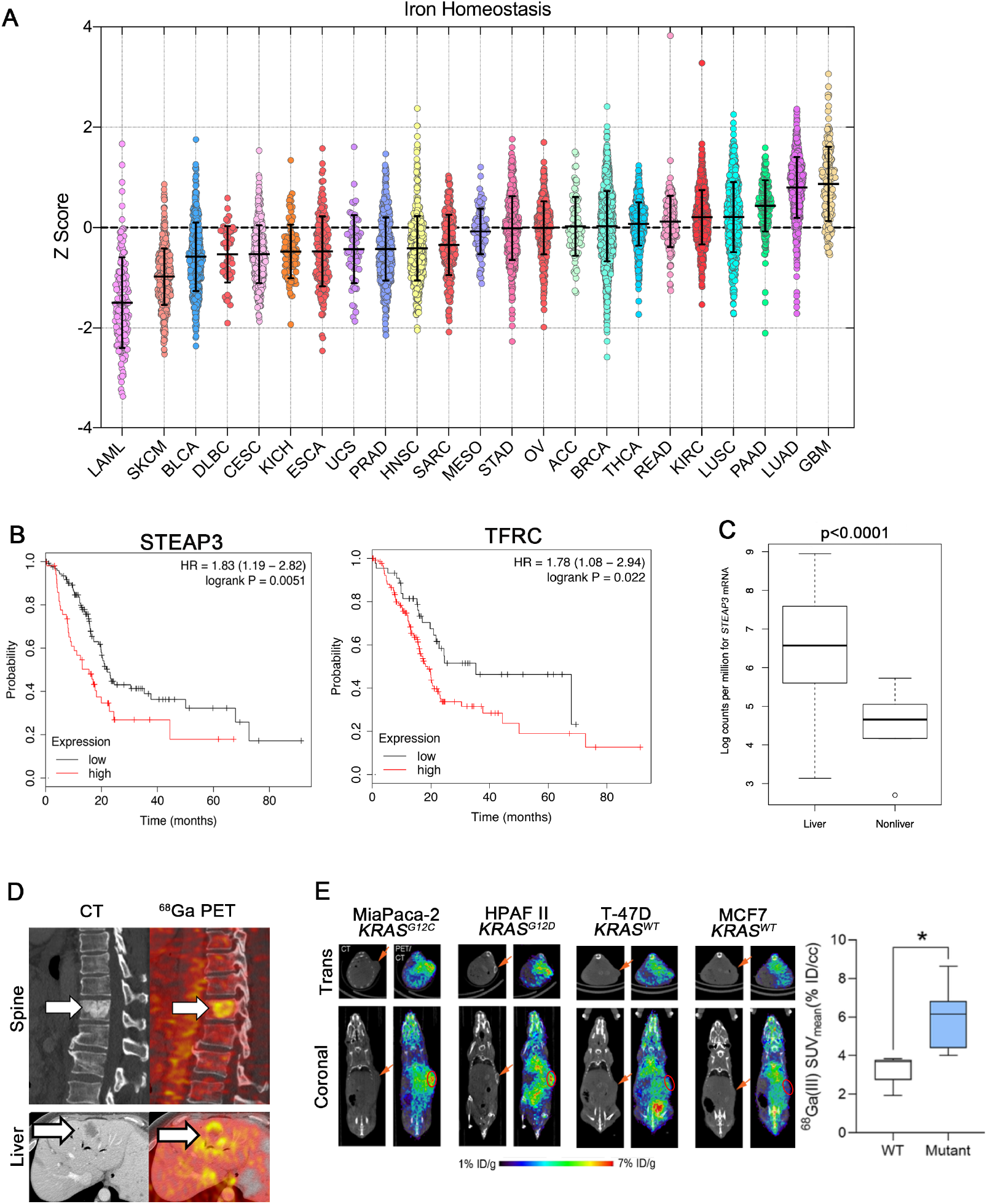
A ferrous iron signature is prognostic in pancreatic adenocarcinoma (A) Pan Cancer analysis of Iron Homeostasis. Z-sores are plotted for each of 23 tumor types. Medians for each tumor type are ranked from lowest to highest (R→L). Error bars represent mean ± SEM. (B) PAAD patients were binarized into high (upper half) or low (lower half) of mRNA expression of *STEAP3* and *TFRC*. Survival in months was estimated by the method of Kaplan and Meir. HR= Hazard Ration for Overall Survival. (C) Boxplot showing Median *STEAP3* expression of in hepatic vs. other disease sites in PDA patients profiled in (Aguirre et al., 2018) (D) Representative CT(Left) and fused Ga^68^ citrate PET/CT (Right) images demonstrating Ga^68^ citrate avid metastases within the T10 vertebral body and liver from two patients with pancreatic ductal adenocarcinoma. (E) Representative Ga^68^ PET images of mice bearing xenografts with wild type or mutant KRAS. Tumors bearing an oncogenic KRAS mutation have an increased avidity for Ga^68^. Error bars represent mean ± SEM, n = 3 mice/group and analyzed by two sample t test. *p < 0.05.

To better understand these clinical observations, we sought to explore Fe^3+^ and Fe^2+^ avidity in xenograft mouse models. We observed enhanced uptake of Gallium-68 in mutant KRAS (*KRAS^m^*) driven PDA xenografts as compared to breast cancer xenografts with wt KRAS (**Figure 1E**). To assess levels of reduced iron in the same tumors, we used ^18^F-TRX, a recently described radiotracer capable of detecting oxidation-state specific for Fe^2^ (Muir et al., 2019). We found that the mKRAS-driven PDA tumors were enriched in Fe^2+^ as compared to the wt KRAS tumors, mirroring the findings with ^68^Ga (**Figure S1A**). Together these findings suggest that increased iron uptake and iron reduction is common in PDA and associated with poor prognosis in patients.

### Oncogenic KRAS Drives Accumulation of Labile Ferrous Iron

We next investigated the mechanism(s) by which oncogenic *KRAS* drives Fe^2+^ accumulation. To temporally control *KRAS^G12D^* expression in primary cells, we isolated primary dermal fibroblasts from a Lox–Stop–Lox *KRAS* (*LSL-KRAS^G12D^*) conditional mouse model, in which the expression of *KRAS^G12D^* is controlled by removal of a Cre recombinase-sensitive transcriptional Stop element. We achieved removal of the Stop element from the *LSL-KRAS^G12D^* allele by the use of an adenovirus expressing Cre recombinase (AdenoCre). We collected protein from primary fibroblasts at 24hr, 48hr and 72hr after AdenoCre infection and observed robust expression of KRAS^G12D^ as well as increased MAPK signaling, as demonstrated by phospho-ERK induction (**Figure 2A**). Intracellular labile Fe^2+^ also increased, as measured with the Fe^2+^-selective florescent dye SiRhoNox (Hirayama et al., 2019) (**Figure 2B,C**). Proliferation slowed upon *KRAS^G12D^* expression and p21^CIP^ levels increased, consistent with oncogene induced senescence (Serrano et al., 1997) (**Figure S1B-C**). To extend our findings we also isolated primary pancreatic ductal cells (Reichert et al., 2013) from Lox–Stop–Lox *Kras* (*Kras^LSL_G12D^*); *Trp53^flox/flox^* and again activated expression of Kras^G12D^ with AdenoCre transduction. We again observed increased Fe^2+^ levels at 72h after AdenoCre (**Figure 2A,D**). Similarly, increased intracellular ferrous iron was observed in both HPDE cells (immortalized human pancreatic duct epithelial cells) and HBEC cells (immortalized human bronchial epithelial cells) after expression of *KRAS^G12D^* by lentiviral transduction (**Figure 2E**). Evaluation of a larger panel of human malignant cell lines from PDA and breast cancer confirmed that cells harboring mutant *KRAS* had consistently higher levels of total intracellular ferrous iron (**Figure 2F**). These results demonstrate early and sustained increases in Fe^2+^ as a consistent effect of oncogenic KRAS expression and distinctly uncouple Fe^2+^ accumulation from cellular proliferation in primary cells which undergo a Fe^2+^-rich replicative arrest upon acute exposure to high levels of KRAS^G12D^ signaling.

**Figure 2.**
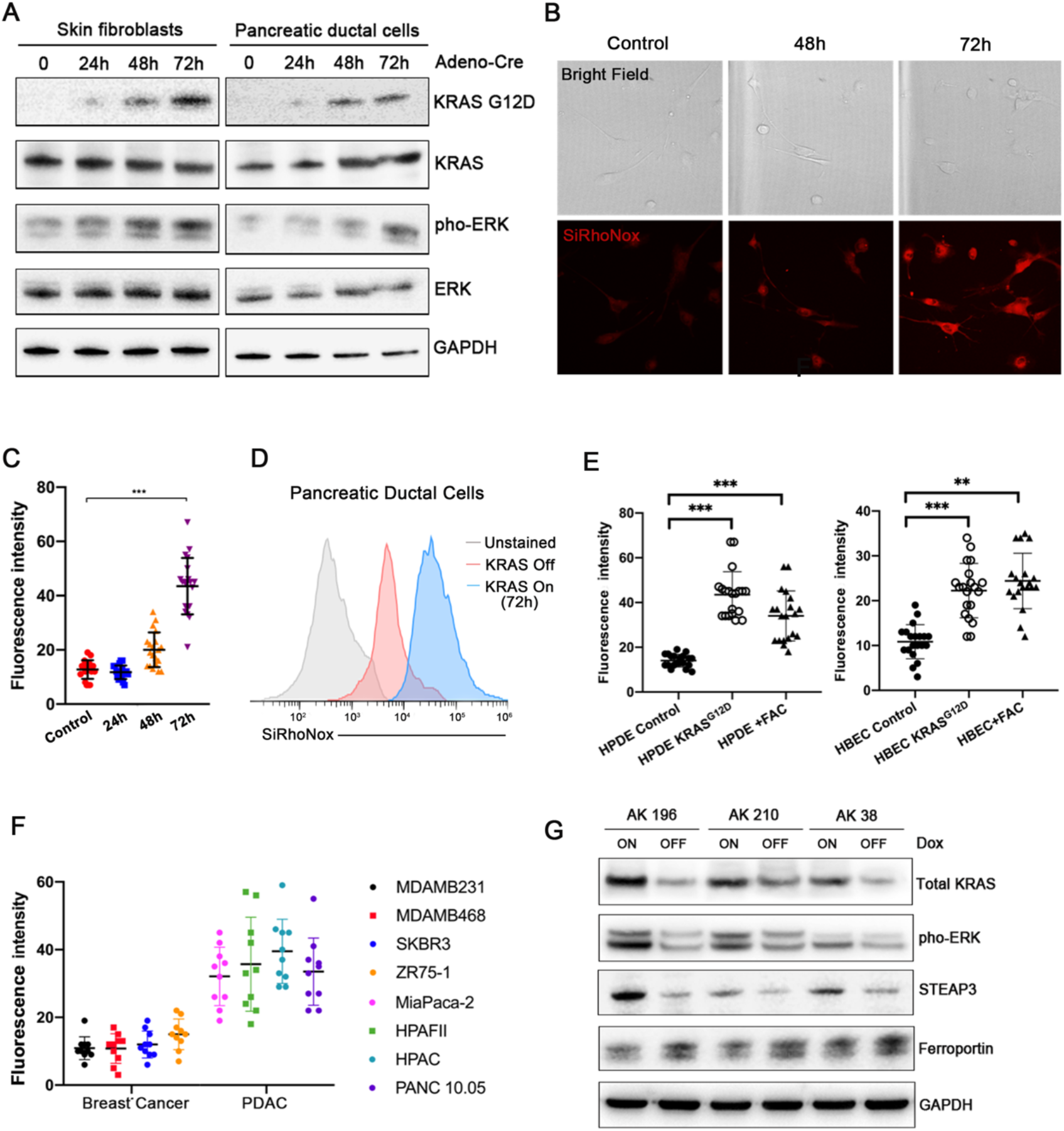
Oncogenic KRAS drives an accumulation of labile ferrous iron (A) Western blot for KRAS^G12D^, KRAS, phosphorylated ERK and total ERK in *LSL-KRAS^G12D^*, *p53^f/f^* mouse pancreatic ductal cells and *LSL-KRAS^G12D^* mouse fibroblasts before and after treatment with AdenoCre at 24h, 48h and 72h. (B-C) Representative images of bright field and IF staining for SiRhoNox in *LSL-KRAS^G12D^* mouse fibroblasts. Quantifications of the fluorescence intensity per cell are plotted. (D) Representative histogram assessing staining for SiRhoNox by flow cytometry in *LSL-KRAS^G12D^*, *p53^f/f^* mouse pancreatic ductal cells before and after the treatment with AdenoCre at 72h. (E) Quantifications of the fluorescence intensity stained with SiRhoNox in HPDE, HPDE KRAS^G12D^ cells and HPDE cells treated with ferric ammonium citrate (FAC). And quantifications of the fluorescence intensity stained with SiRhoNox in HBEC, HBEC KRAS^G12D^ cells and HBEC cells treated with ferric ammonium citrate (FAC). (F) Quantifications of the fluorescence intensity stained with SiRhoNox in various breast and pancreatic cancer cell lines. (G) Western blot for Kras^G12D^, total Kras, phosphorylated Erk1/2 and total Erk1/2 in iKRAS cells treated with or without doxycycline. Error bars represent mean ± SEM, n=10cells/group in B, n=20cells/group in C-E and n=3 replicates in G and analyzed by two sample t test. **p < 0.01, ***p < 0.001.

To further elucidate the effect of oncogenic KRAS on iron metabolism and transport, we deployed previously reported genetically engineered mouse PDA cancer cell lines (designated *iKras^G12D^*) in which the expression of *Kras^G12D^* is dependent upon continuous doxycycline exposure (Ying et al., 2012). Partial extinction of Kras protein and phospho-ERK followed doxycycline withdrawal within 48hr. Three *iKras* lines: AK38, AK196 and AK210, all exhibited significant decreases in *STEAP3* expression and elevation of *FPN* (Ferroportin, an iron exporter) expression (**Figure 2G, Figure S1D**) after Dox withdrawal. Expression of oncogenic *KRAS* resulted in similar effects on *STEAP3* and *FPN* expression in three non-transformed cell lines HPDE, AALE and MEFs (**Figure S1E**). These data suggest that oncogenic KRAS is both necessary and sufficient to maintain an elevated ferrous iron pool, at least in part by increased expression of ferrireductase and reduced iron efflux.

### A Ferrous Iron–Activatable Drug Conjugate (FeADC) Targets Elevated Ferrous Iron

*KRAS* is the most commonly mutated oncogene in human cancer but has proven difficult to pharmacologically target. RAF-MEK-MAPK is the major KRAS effector arm in PDA (Collisson et al., 2012) acute myeloid leukemia (Burgess et al., 2017) and lung adenocarcinoma (Karreth et al., 2011). Fe^2+^-dependent pharmacology (wherein a drug is activated only in the presence of Ferrous iron) has been established in the antimalarial artemisinin but is mostly unexplored outside of anti-protozoal therapies. Since we found elevated Fe^2+^ levels to be driven by oncogenic KRAS, we hypothesized that KRAS-driven PDA tumor cells might be selectively targeted with a Ferrous Iron–Activatable Drug Conjugate (FeADC) based on 1,2,4-trioxolane (TRX) chemistry (Fontaine et al., 2014). To determine whether an FeADC might be effective in PDA, we probed PDA cells with the Fe^2+^ probe TRX-PURO, a Fe^2+^-activatable conjugate of the aminonucleoside puromycin (Spangler et al., 2016). Puromycin is released from TRX-PURO in cells via reaction with Fe^2+^ (**Figure 3A**), the degree of PURO release occurring in cells is quantifiable with a puromycin antibody. We treated primary skin fibroblasts harboring cre-inducible *KRAS^G12D^* with TRX-PURO after synchronously inducing high levels of oncogenic *KRAS^G12D^*. We observed a time-dependent increase in puromycin signal out to 72hr post-infection (**Figure S1F**), mirroring our findings with SiRhoNox (**Figure 2B-D**), and indicating the efficient Fe^2+^-dependent activation of TRX-PURO in cells expressing oncogenic KRAS.

**Figure 3.**
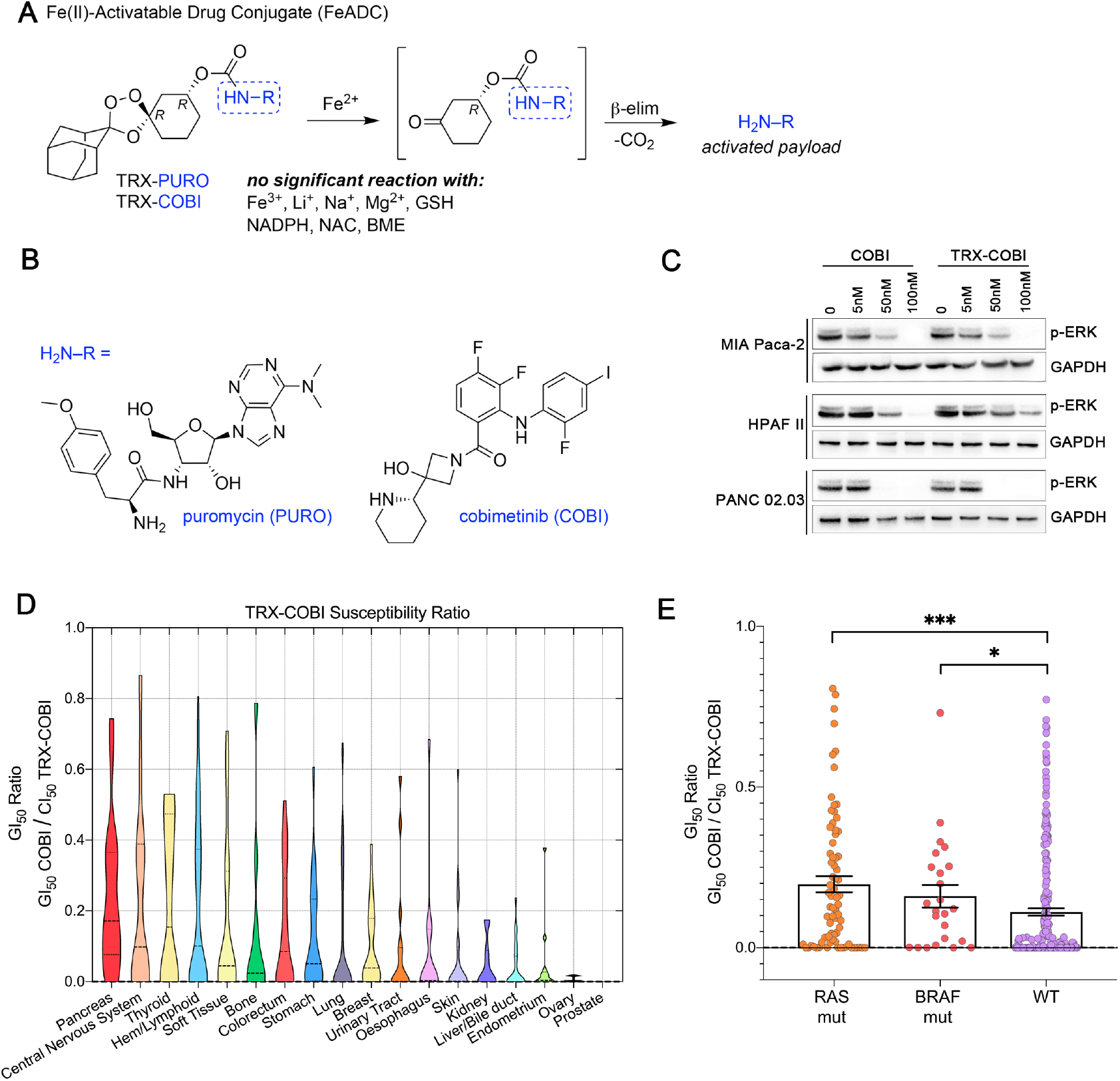
Ferrous Iron–Activatable Drug Conjugate (FeADC) targets elevated ferrous iron in PDA (A) Structure and mechanism of FeADC activation by ferrous iron sources. (B) Chemical structure of FeADC payloads puromycin and cobimetinib (COBI). (C) Western blot for phosphorylated ERK in COBI or TRX-COBI treated MIAPaca-2, HPAFII and PANC 02.03 cells at indicated doses. (D) A large panel of cancer cells were dosed with COBI and TRX-COBI for 5 days in PRISM screen. The GI50 of TRX-COBI for each cell was normalized by that of the GI50 to parent COBI to generate a TRX susceptibility ratio for each cell line. (E) *RAS*, *BRAF* mutation status and GI_50_ ratios from PRISM screen.

Having established that mutant KRAS elevates intracellular Fe^2+^ as measured with both SiRhoNox and TRX-PURO, we sought to inhibit the RAF-MEK-ERK pathway in PDA with a new FeADC based on TRX chemistry. Targeting MEK with allosteric inhibitors of MEK1/2 has shown clinical benefit but the approach suffers from on-target toxicities that are dose limiting in the eye, skin, gut and other organs. Clinical experience has shown sustainable dosing of these agents is typically only ~25% of the FDA-approved dose (Saunders et al., 2020) severely hampering the dose intensity achievable in the tumor cell, and ultimately limiting clinical efficacy. To model MEK inhibitor toxicity in animals, we administered the MEK inhibitor binimetinib to healthy mice for three weeks. We observed thinning of the epidermal keratinocyte layer in tail skin (**Figure S2A**) confirming the conserved, essential role for MEK1/2 in the cellular epidermal layer(Azan et al., 2017; Danilenko et al., 2016). We hypothesized that a FeADC targeting MEK1/2 would be efficiently activated by Fe^2+^ in mKRAS tumors, but not in healthily mouse epidermis, where normal iron homeostasis is intact. We synthesized FeADCs bearing the FDA-approved allosteric MEK inhibitor cobimetinib (COBI) and based on either enantiomeric form of the Fe^2+^-targeting TRX moiety, yielding the novel agents (*R*,*R*)-TRX-COBI and (*S*,*S*)-TRX-COBI (**Figure 3B**). Incubation of these FeADCs in mouse and human liver microsomes revealed somewhat superior metabolic stability of (*R*,*R*)-TRX-COBI (**Supplementary Information**) and so this form was used in all subsequent studies (henceforth denoted simply TRX-COBI). We next evaluated the pharmacokinetic properties of TRX-COBI in NSG mice following a single 15 mg/kg dose by the IP route. We observed a favorable exposure profile, with an elimination half-life of ~6.9 hrs, C_max_ = 1421 ng/mL, and AUC_inf_ = 6462 h*ng/mL (full parameters provided in **Supplementary Information**). Significantly, the extent of free COBI released from TRX-COBI in this study was minimal, ca. ~2% of the total dose by C_max_ and AUC_inf_. Overall, the pharmacokinetic properties of TRX-COBI were similar to that reported previously for COBI in Nu/Nu mice (Choo et al., 2012), suggesting TRX-COBI as a suitable FeADC prototype for *in vivo* study.

We compared the growth inhibitory effects of TRX-COBI and COBI across a panel of human PDA cells harboring m*KRA*S and found that TRX-COBI was efficiently activated, as evidenced by similar phospho-ERK reductions at equimolar exposure of COBI (**Figure 3C**). We further explored the relationship of TRX deconjugation efficiency with cell type and genotype using the PRISM prlaform (Yu et al., 2016). We screened COBI and TRX-COBI in 8-point dose response across 750 cancer cell lines spanning 18 tumor types. Briefly, cell lines were treated with inhibitors for 5 days, then the relative abundance of each cell line was determined by high-throughput sequencing of barcodes and generated growth inhibitory doses (GI50) of TRX-COBI and COBI for each cell line. We then calculated a TRX-COBI susceptibility ratio (i.e. GI50_COBI_ / GI50_TRX-COBI_) for 520 out of 750 cell lines passing criteria needed for index creation (Supplementary information).

PDA cells exhibited the highest mean TRX-COBI GI50 ratio across 18 type of tumor cells (**Figure 3D**). Additionally, *BRAF* and *RAS* mutated cancer cell lines also had significantly higher TRX-COBI GI50 Ratios than wild-type RAS lines (P < 0.001; **Figure 3E**), indicating more efficient activation of FeADCs in MAPK-driven tumor cells, irrespective of histology.

To assess the known modifiers of intracellular Fe^2+^ in cells we depleted *STEAP3* in MiaPaca2 and Capan1 cells with shRNA which led to (~50%) decreases in intracellular Fe^2+^ levels (**Figure S2C**). As expected, *STEAP3*-depletion decreased conversion of TRX-COBI to COBI as measured by phospho-ERK/ERK ratios (**Figure S2D-E**). Together these data indicated that a prototypical MEK inhibitor FeADC is effective in KRAS-driven PDA cells and that FeADC activation depends, at least in part, on the cellular ferroreductive machinery shown above to be under control of oncogenic KRAS.

### An FeADC of Cobimetinib Exhibits Potent Anti-Tumor Activity *in vivo*

We next evaluated the FeADC approach to MEK1/2 inhibition *in vivo*. We orthotopically implanted mouse *Kras^LSL_G12D^*, *Trp53^f/+,^ Pdx1^Cre^* (KPC) cells into syngeneic FVB mice and treated with equimolar doses of either TRX-COBI or COBI. Both COBI and TRX-COBI were comparably tumor growth inhibitory (**Figure 4A-B**) and showed equivalent suppression of phospho-ERK, at equimolar doses in this orthotopic PDA model (**Figure 4G-H**). We next evaluated the effects of COBI and TRX-COBI on overall survival in a widely used autochthonous *Kras^LSL-G12D/+^; Trp53^flox/flox,^ (KP)* mouse lung cancer model (Jackson et al., 2001). We initiated treatment with equimolar doses of COBI, TRX-COBI or vehicle 8 weeks after adenoviral tumor induction for 60 days. Mice receiving either COBI or TRX-COBI (equimolar doses) had fewer lung lesions (as evidenced by LSL-Tomato florescence) and showed prolonged overall survival compared to vehicle treated mice (**Figure 4C-D**). We further examined the therapeutic efficacy of TRX-COBI in patient-derived xenograft (PDX) models. We used two *KRAS^G12D^*-driven PDA patient derived xenograft (PDX) models and a *KRAS^G12C^*-driven non-small cell lung cancer (NSCLC) PDX. Both COBI and TRX-COBI each significantly inhibited tumor growth relative to vehicle-treated tumors (**Figure 4E-F**). In addition, tumor lysates as well as tumor immunohistochemistry from PDA 260 showed that TRX-COBI reduced pERK as effectively as parental COBI (**Figure 4G-H**). Collectively, these data demonstrate equivalent *in vivo* anti-tumor activity with equimolar doses of TRX-COBI or COBI in multiple relevant *KRAS*-driven models of lung and pancreatic adenocarcinoma *in vivo*.

**Figure 4.**
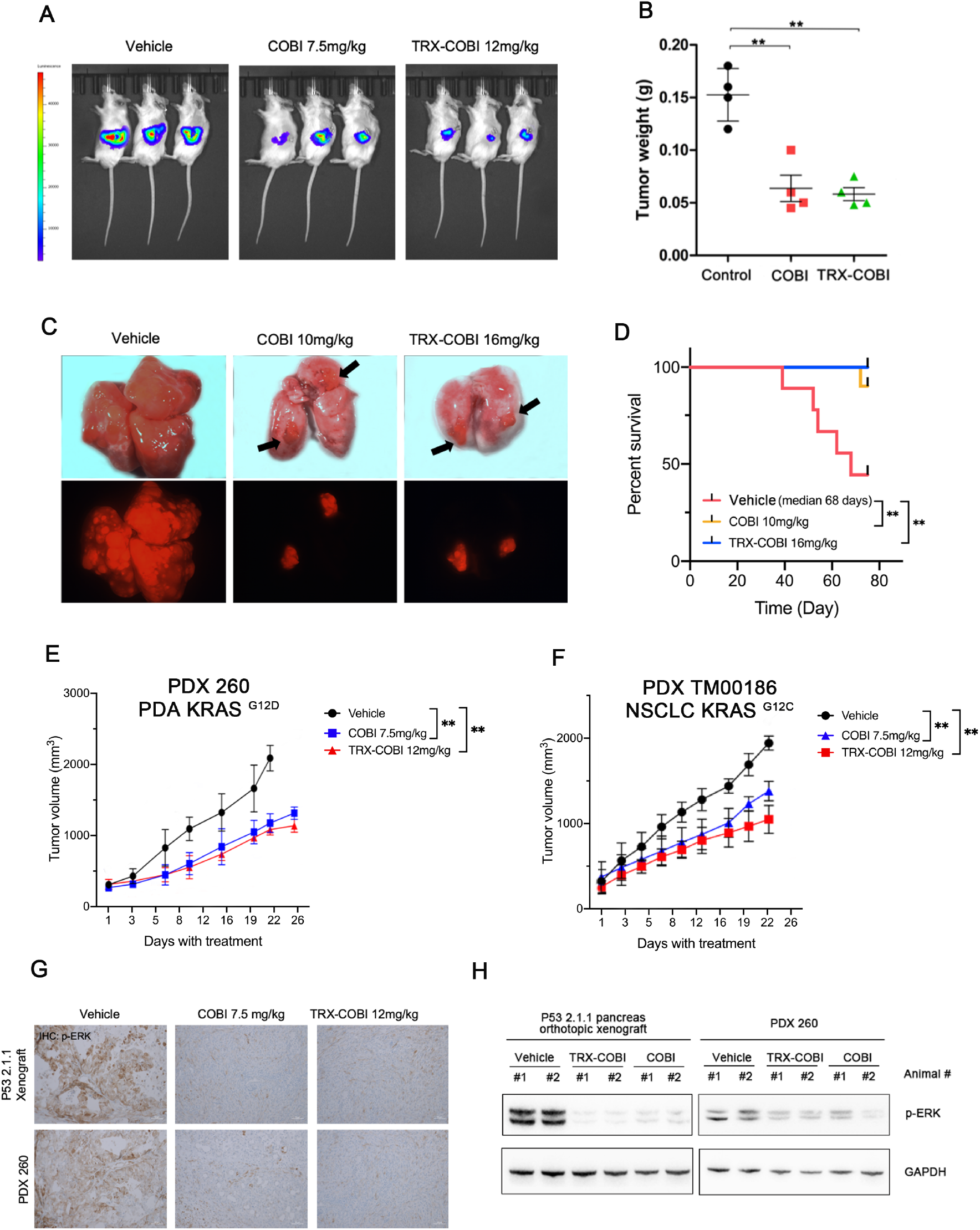
An FeADC of Cobimetinib exhibits potent anti-tumor activity *in vivo* (A) Representative bioluminescence images for the effects of vehicle and the equimolar dose of COBI and TRX-COBI on p53 2.1.1-fLuc orthotopic pancreas tumor xenografts. (B) Tumor weights at the end point from A. Error bars represent mean ± SEM, n=4-5mice/group and analyzed by two-sample t-test. **p < 0.01. (C) Representative gross lung images in *KP* mouse model. Vehicle, equimolar dose of COBI and TRX-COBI treatment started 8 weeks after the adenoviral induction and continued for 60 days. (D) Overall survival of (C) analyzed by logrank test, n=8-10mice/group, **p < 0.01. (E-F) Tumor volume changes (mean ± SEM; error bars) of KRAS mutant PDA and NSCLC PDX model, n=8-10mice/group. Data were analyzed by one-way ANOVA **p < 0.01. (G) IHC of xenografts from A and E (10X) stained with phosphorylated ERK. (H) Western blot for phosphorylated ERK of tumors resected at the end of treatment in A and E.

### FeADC Approach Mitigates MEK Inhibitor On-Target Off-Tumor Toxicities

We next asked whether the FeADC approach might mitigate on-target, off-tissue toxicities of systemic MEK inhibition, reasoning that TRX-COBI should remain inactive (TRX-bound) in normal tissues (skin, retina) that are adversely impacted by systemic MEK inhibition in cancer patients. We used retinal cells and keratinocytes as experimentally tractable models of two tissues of known MEK inhibitor toxicity. We found that human retinal pigment epithelia cells (RPE-1) and keratinocytes (HaCaT) were about 10-fold less sensitive than PDA cells to TRX-COBI, as evidenced by intact pERK levels in the off-target cells at drug concentrations sufficient to completely abolish pERK in PDA cells. In stark contrast, COBI more potently inhibited pERK in off-target keratinocytes than in PDA cells (**Figure 5A**). These results suggest that the FeADC approach can realize improved therapeutic index in mKRAS-driven cancers.

**Figure 5.**
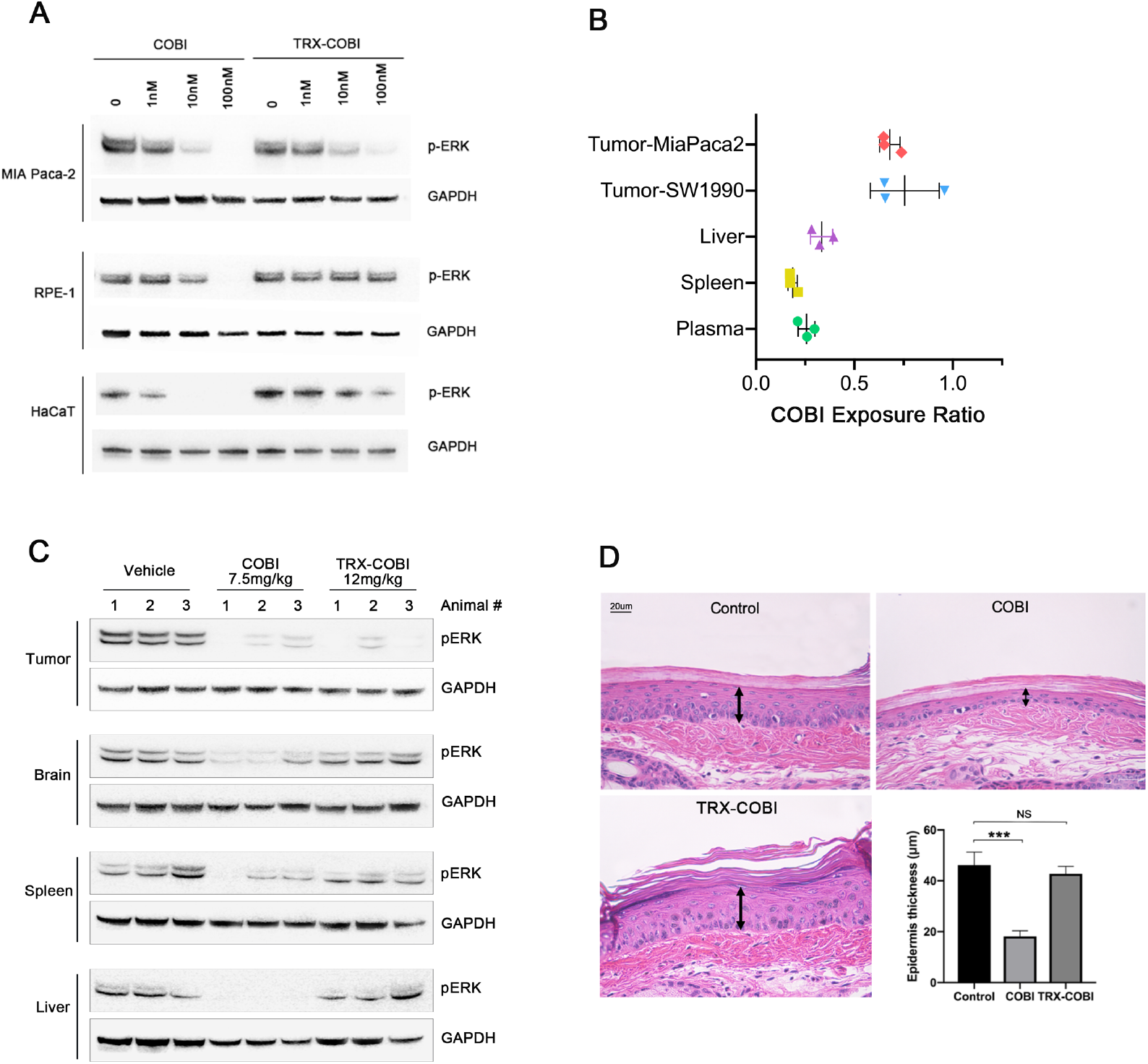
FeADC approach mitigates MEK inhibitor Off-Tumor/On-Target toxicities (A) Western blot for phosphorylated ERK of MiaPaca-2, RPE-1 and HaCaT cells treated with COBI and TRX-COBI at indicated dose. (B) COBI exposure ratios in tumors, plasma, spleen and liver after treatment for 7h. Data was determined by LCMS/MS assay. Error bars represent mean ± SEM. (C) Western blot for phosphorylated ERK of tumor xenografts, brain, spleen and liver from NSG mice after treatment with vehicle, the equimolar dose of COBI and TRX-COBI for 8h. (D) Representative images of HE (40X) show the epidermal layer of mouse tail skin after the treatment with vehicle, the equimolar dose of COBI and TRX-COBI for 20days. Error bars represent mean ± SEM, n=3 mice/group and analyzed by two sample t test. ***p < 0.001.

Next, we explored the tissue-selective activation of FeADCs in animals. We administered a single, equimolar dose of TRX-COBI, COBI or vehicle, to mice bearing subcutaneous SW1990 and MiaPaca2 tumors in each flank followed 7 hours later by collection of tissue from brain, liver, spleen, tail skin, plasma and tumor. We calculated a COBI exposure ratio by dividing the concentration of free COBI in the tumor/tissues of mice receiving TRX-COBI with those receiving COBI directly, i.e. COBI exposure ratio = [COBI]_TRX-COBI treated_ / [COBI]_COBI treated_. We observed higher COBI exposure ratios in both SW1990 and MiaPaca2 tumors compared to plasma, liver and spleen (**Figure 5B**). Phospho-ERK was significantly and comparably reduced in tumor and nontumor tissues of the COBI-treated mice, unsurprisingly, reflecting equivalent MEK inhibition in tumor and healthy tissues. In contrast, phospho-ERK was extinguished in the tumors of in TRX-COBI treated mice to an equivalent degree as in COBI-treated mice, however phospho-ERK was spared (similar to vehicle control) in the brain, spleen, liver, and tail skin (**Figure 5C and Figure S3A**). This was consistent with the higher COBI exposure ratios observed in tumor vs. normal tissues and indicates that TRX-COBI remains inactive in normal tissues from tumor-bearing animals while still extinguishing KRAS-induced MAPK signaling in the tumor itself. To model the cumulative toxicities of chronic, systemic MEK inhibition, we collected tail skin samples from mice dosed with equimolar amounts of TRX-COBI, COBI or vehicle for 20 days and measured the cellular epidermal layer. We consistently found reduction of the epidermal layer in mice dosed with COBI (**Figure 5D and Figure S3B**) but not in TRX-COBI dosed animals. Together, these findings show that many of the MEK1/2-dependent (on target) extra-tumoral toxicities (Scholl et al., 2009),(Balagula et al., 2011) of MEK inhibition are substantially mitigated by the FeADC approach, which shows promise to improve the poor therapeutic index of RAS-RAF-MAPK blockade in the clinic.

### MEK Inhibition with FeADC Enables Dose Intensity in Combination Therapies

The inherently low therapeutic index of MEK inhibitors has hindered development of MEKi-containing therapeutic combinations. Our encouraging data with TRX-COBI monotherapy argued for improved tolerance of combination treatment schemes anchored upon MEK inhibition combined with inhibition of other RAS effectors or activators. Inhibition of SHP2 can sensitize many cycling dependent *KRAS*-mutant or *KRAS*-amplified cancers to MEK inhibitors (Ruess et al., 2018). While such vertical blockade holds tremendous promise, combination therapy of SHP2 and MEK inhibitors negatively effects tolerability in mice (Fedele et al., 2018). Indeed, we found the combinations of tolerable monotherapy doses of COBI and the SHP2 inhibitor SHP099 (Novartis) induced study-terminating body weight loss from baseline within three weeks when combined in non-tumor bearing NSG mice (**Figure S4A**).

To assess the potential of FeADC strategies to improve the toxicity profile of vertical MAPK pathway blockade, we first assessed cytotoxicity of combined MEK+SHP2 dual inhibition in PDA (MiaPaca2) vs. normal (RPE-1 and HaCaT) cells. Treatment of non-malignant RPE-1 and HaCaT cells with increasing concentrations of COBI enhanced cell growth inhibition of SHP2 inhibitor (SHP099, 10uM) while equimolar combinations of TRX-COBI/SHP2i were less toxic to these normal cells (**Figure 6A-B**). In contrast, *KRAS^G12C^* MiaPaca-2 cells showed equivalent sensitivity to the two combinations (**Figure 6C**), indicating a potentially improved safety profile with and comparable efficacy of TRX-COBI in MEK inhibitor-based combination therapies.

**Figure 6.**
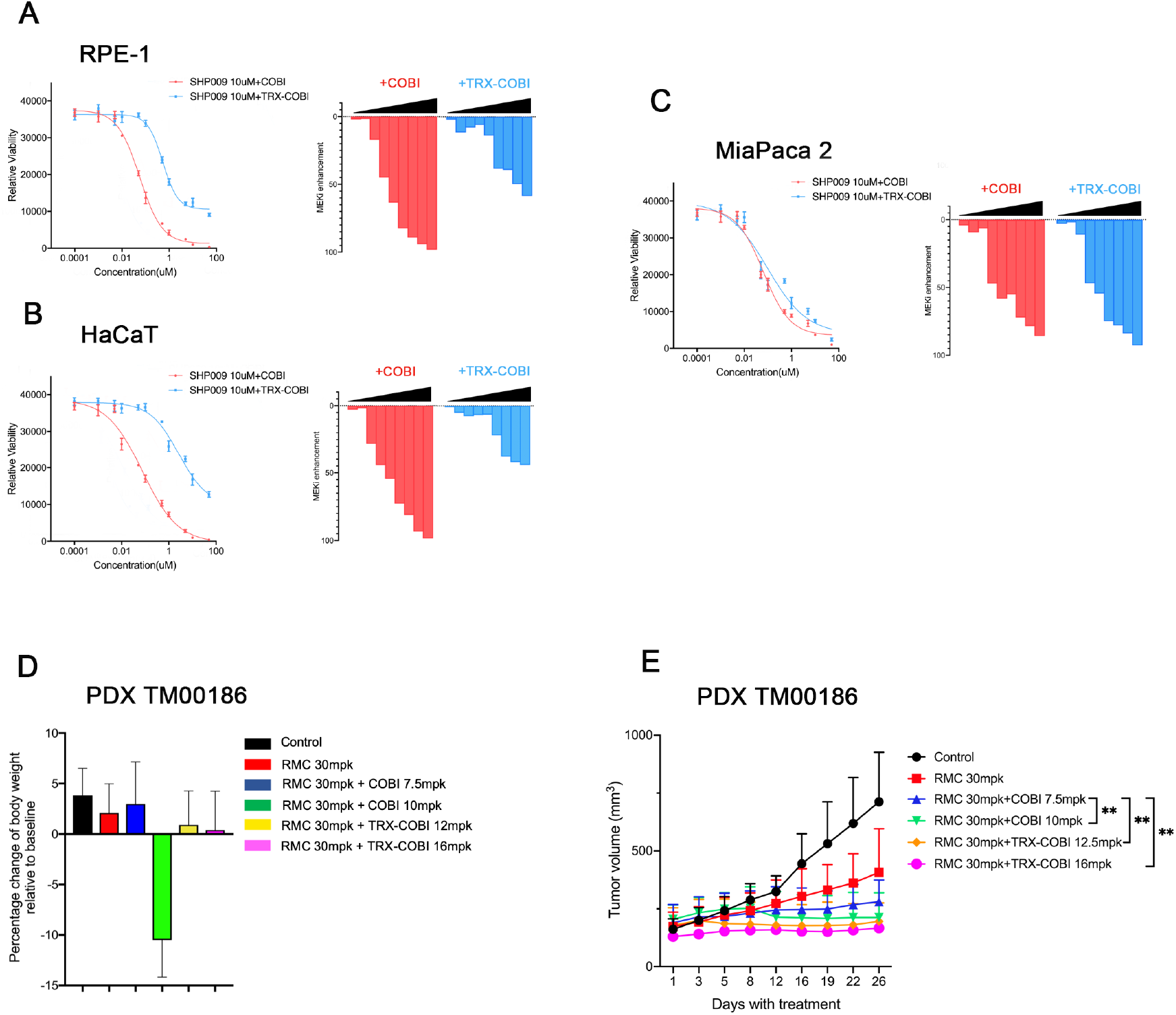
MEK inhibition with FeADC enables dose intensity in combination therapies (A-C) RPE-1, HaCaT and MiaPaca2 cells were treated with increasing concentrations of COBI or TRX-COBI in the presence of 10uM SHP009 for 72 hours, and viability was determined. The enhancement of growth inhibition by the addition of COBI or TRX-COBI to SHP009 was determined by normalizing the viability after combination treatment to SHP009 for each dose point (0 = no effect; 100% = complete cell killing). (D) Mouse body weight changes normalized to the baseline after the treatment of vehicle, 30 mg/kg RMC-4550 monotherapy, or combinations comprising 30 mg/kg of RMC-4550 combined with COBI at either 7.5 mg/kg or 10 mg/kg, or RMC-4550 with comparable (equimolar) 12.5 mg/kg or 16 mg/kg doses of TRX-COBI. Error bars represent mean ± SEM, n=6mice/group. (E) Tumor volume changes of PDX TM00186 tumors after the treatments in D. Error bars represent mean ± SEM, n=6mice/group and analyzed by one-way ANOVA **p < 0.01.

We next compared the efficacy and tolerability of TRX-COBI to COBI when combined with the SHP2 inhibitor RMC-4550 (Revolution Medicines) in *KRAS^G12C^* NSCLC (TM00186) PDX model. Six separate groups of mice were administered vehicle, 30 mg/kg RMC-4550 monotherapy, or combinations comprising 30 mg/kg of RMC-4550 combined with COBI at either 7.5 mg/kg or 10 mg/kg, or RMC-4550 with comparable (equimolar) 12.5 mg/kg or 16 mg/kg doses of TRX-COBI. Of note, mice receiving the combination of COBI 10mg/kg plus RMC 30mg/kg lost weight (~10%) (**Figure 6D**) and exhibited lassitude. Tumor and spleen lysates both showed profound inhibition of the expression of phospho-ERK (**Figure S4B-C**). Remarkably, the equivalent combination of TRX-COBI 16 mg/kg with RMC 30mg/kg induced the most pronounced tumor growth inhibition without measurable weight loss (**Figure 6E**) and showed minor effects on the inhibition of phospho-ERK in the spleens (**Figure S4B-C**). Together, these data indicate that FeADC-based MEK inhibition combined with SHP2 blockade exhibits superior tolerability and improved efficacy in KRAS driven tumors.

## Discussion

Iron in the form of enzyme cofactors powers energy generation and macromolecule synthesis essential for cell proliferation and survival. Consistent with observations in breast cancer (Miller et al., 2011; Pinnix et al., 2010), prostate cancer (Tesfay et al., 2015), and glioblastoma (Schonberg et al., 2015), we have now confirmed an iron avid phenotype in PDA. More specifically, it is an elevated pool of iron in the Fe^2+^ state that characterizes PDA and likely also in other cancers. Ferroaddiction comes with new liabilities (Dixon et al., 2012), since labile Fe^2+^ promotes intracellular Fenton chemistry, in turn resulting in elevated intracellular reactive oxygen species (ROS) (Dizdaroglu and Jaruga, 2012; Inoue and Kawanishi, 1987) and ferroptosis-associated lipid peroxidation that in turn invokes a constitutive dependence on ROS-detoxifying enzymes like GPX4 (Forcina and Dixon, 2019; Liu et al., 2018) and/or FSP1 (Bersuker et al., 2019; Doll et al., 2019). Our findings in in KRAS-driven cells and tumors provide additional evidence linking oncogenic KRAS to elevated Fe^2+^ and a vulnerability to ferroptosis (Dixon et al., 2012).

Iron homeostasis is tightly controlled at the level of the cell and organism and the dysregulation of iron regulatory genes in cancer biology has been recognized for some time (Torti and Torti, 2013b). The mechanisms by which specific oncogenes drive iron utilization to sustain tumor growth is unclear. Crucially, the oxidation-state specific study of iron has been impossible prior to the recent advent of reactivity-based chemical probes like SiRhoNox and TRX-PURO that are highly selective for the much less abundant ferrous (Fe2+) ion. Here we show that KRAS signaling is necessary and sufficient to empower Fe^2+^ demand in PDA cells. In particular, oncogenic KRAS exerts control over key iron transporters, exporters, and the lysosomal ferrireductase, which collectively serve to maintain an expanded pool of bioavailable Fe^2+^. Ras oncogenes have been shown to alter nutrient metabolism (Hensley et al., 2016; Perera and Bardeesy, 2015; Ying et al., 2012), and here we provide evidence that oncogenic KRAS signaling also alters iron nutrient metabolism in favor of Fe^2+^ during tumor development.

Most KRAS-mutant cancers depend on sustained expression and signaling of KRAS, thus making it a high-priority therapeutic target (Papke and Der, 2017; Stephen et al., 2014). Unfortunately, development of direct small molecule inhibitors of most mutant K/N/HRAS proteins have been elusive. Moreover, single-agent inhibition of KRAS effectors (e.g. MEK-MAPK pathway) in KRAS-mutant cancers have been disappointing to date (Adjei et al., 2008; Infante et al., 2012; Ko et al., 2016; Zhao and Adjei, 2014). On-target MEK inhibition in healthy tissues dramatically limits the tolerable dose and also the feasibility of MEK inhibitor-based combination therapy. The cellular pathways influenced by dominant acting oncogenes are often essential for homeostatic maintenance of healthy tissues in patients. As a result, potent inhibitors of pathways downstream of mutated oncogenes often have deleterious effects in healthy tissues. This manifests as clinical toxicity. Furthermore, since pathway flux is often greater in cancer cells than in normal cells, the amount of inhibitor required for therapeutic efficacy in the tumor often closely approaches or even exceeds the dose tolerable clinically by healthy tissues.

The iron(II)-mediated Fenton reaction underlies the pharmacology of artemisinins and related endoperoxide antimalarials (Ansari et al., 2013; O’Neill and Posner, 2004). Leveraging this clinically proven chemistry (Fontaine et al., 2014; Mahajan et al., 2012), we introduced the concept of the Ferrous Iron–Activatable Drug Conjugate (FeADC). Having uncovered in the current study an elevation of Fe^2+^ stemming from oncogenic KRAS/MAPK signaling, we synthesized and evaluated TRX-COBI, a novel FeADC designed to target the MAPK pathway in KRAS-driven PDA. Notably, we demonstrate with TRX-COBI tumor selective activation that enables ablation of MAPK signaling in PDA tumor cells and xenografts, while sparing the pathway in normal cells and tissues, including in major organs of iron storage (liver) and targets of MEK inhibitor induced toxicity (skin and CNS). These findings are significant in that they reveal expansion of the bioavailable Fe^2+^ pool in *KRAS* driven, currently poorly treated leathal cancers such as lung and pancreatic adenocarcinoma. We decouple MAPK signaling from elevated Fe^2+^ status in normal and tumor cells, and present a therapeutic strategy that exploits these distinctions to enable more tolerable and efficacious combination therapy targeting the MAPK pathway in KRAS-driven PDA. The discovery of pharmacologically exploitable Ferroaddiction in RAS driven cancers holds promise to improve the treatment of deadly cancers through a practicable and generalizable approach to FeDAC design, development and clinical testing.

---

**Supplementary Figure S1.**
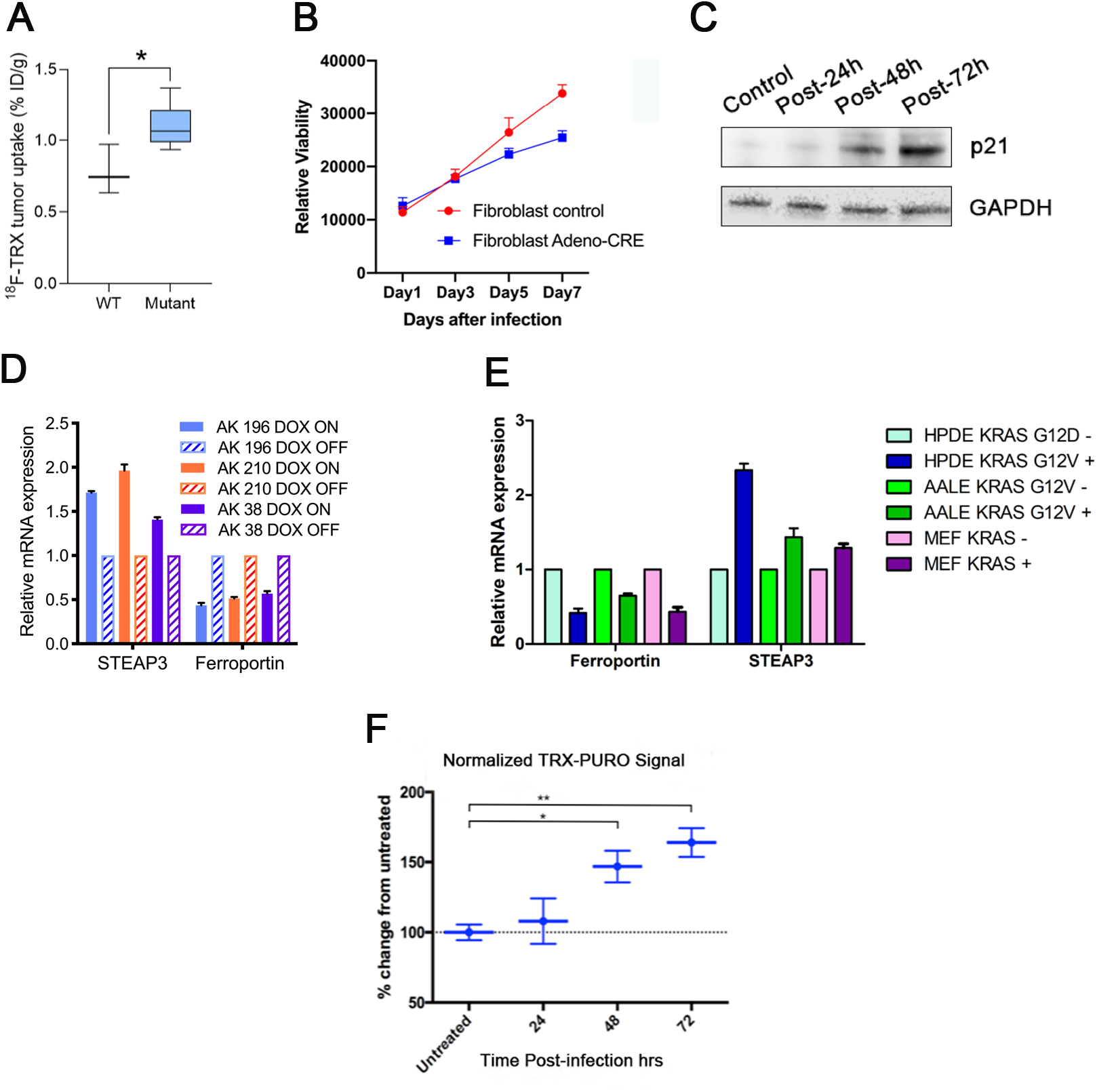
(A) ^18^F-TRX uptake of mice bearing xenografts with wild type or mutant KRAS. Error bars represent mean ± SEM, n = 3mice/group and analyzed by two sample t test. *p < 0.05. (B) Proliferation curve of *LSL-KRAS^G12D^* fibroblasts with or without AdenoCre treatment. (C) West blot for p21 of *LSL-KRAS^G12D^* fibroblasts before and after the treatment with AdenoCre at 24h, 48h and 72h. (D) Quantitative PCR analysis of *Steap3* and *Fpn* (Ferroportin) in iKRAS cells treated with or without doxycycline. (E) Quantative PCR analysis of STEAP3 and Ferroportin expression in indicated cells. (F) Normalized TRX-PURO signals in *LSL-KRAS^G12D^* mouse fibroblasts before and after the treatment with AdenoCre at 24h, 48h and 72h.

**Supplementary Figure S2.**
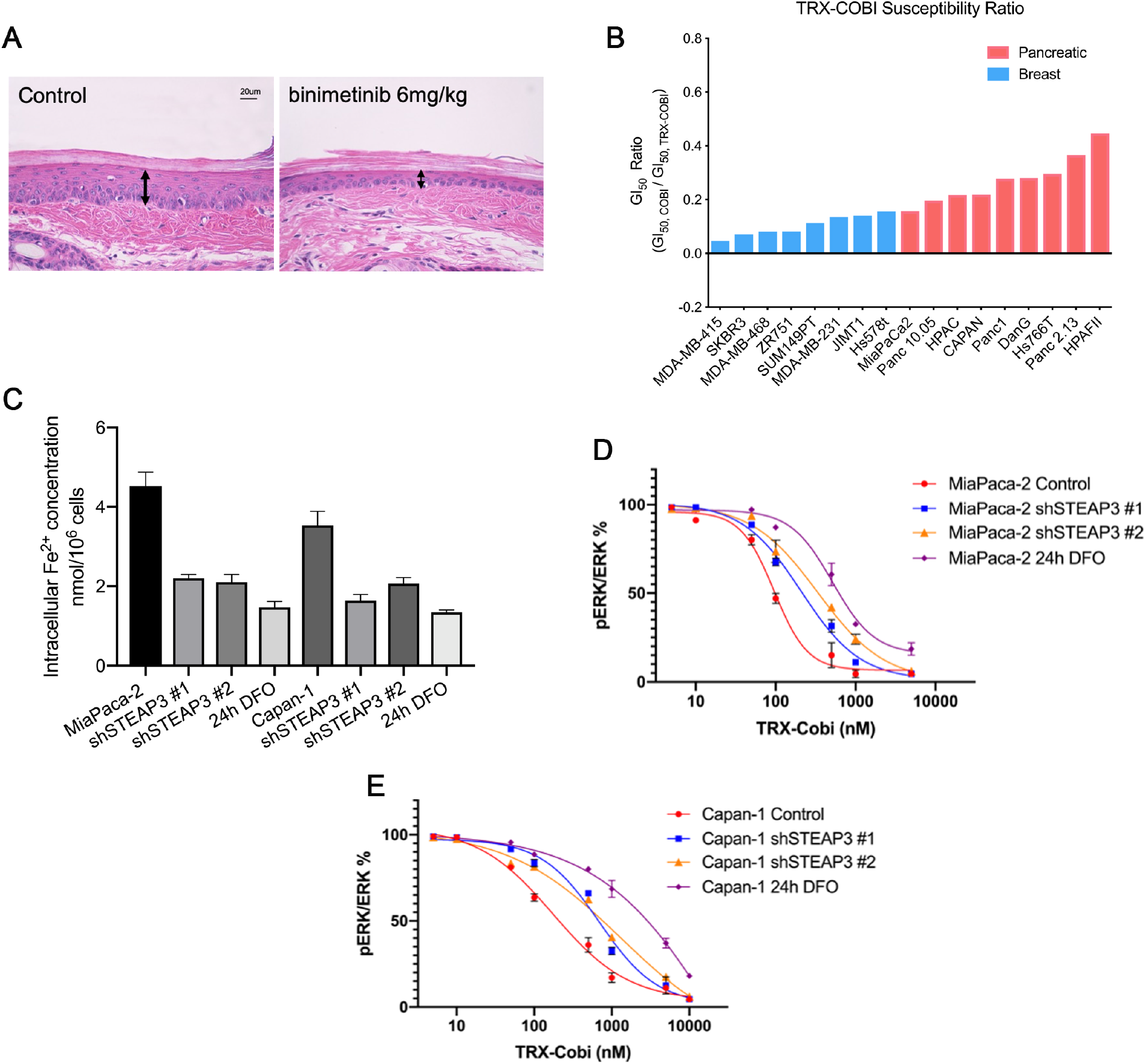
(A) Representative images of HE (40X) show the epidermal layer of mouse tail skin after the treatment with vehicle and Binimetinib after 20days. (B) A panel of pancreatic cancer cells vs breast cancer cells were dosed with COBI and TRX-COBI for 5 days. The GI50 of TRX-COBI for each cell was normalized by that of the GI50 to parent COBI to generate a TRX susceptibility ratio for each cell line. (C) Quantification of intracellular Fe2+ levels in MiaPaca2 and Capan-1 cells with STEAP3 knockdown and iron chelator DFO treatment. (D-E) Phospho-ERK/total ERK levels in t MiaPaca2 and Capan-1 cells with STEAP3 knockdown and prior 24h of iron chelator DFO treatment. Cells were treated with the indicated concentration of TRX-COBI for 2h before lysis. Error bars indicate ±SD.

**Supplementary Figure S3.**
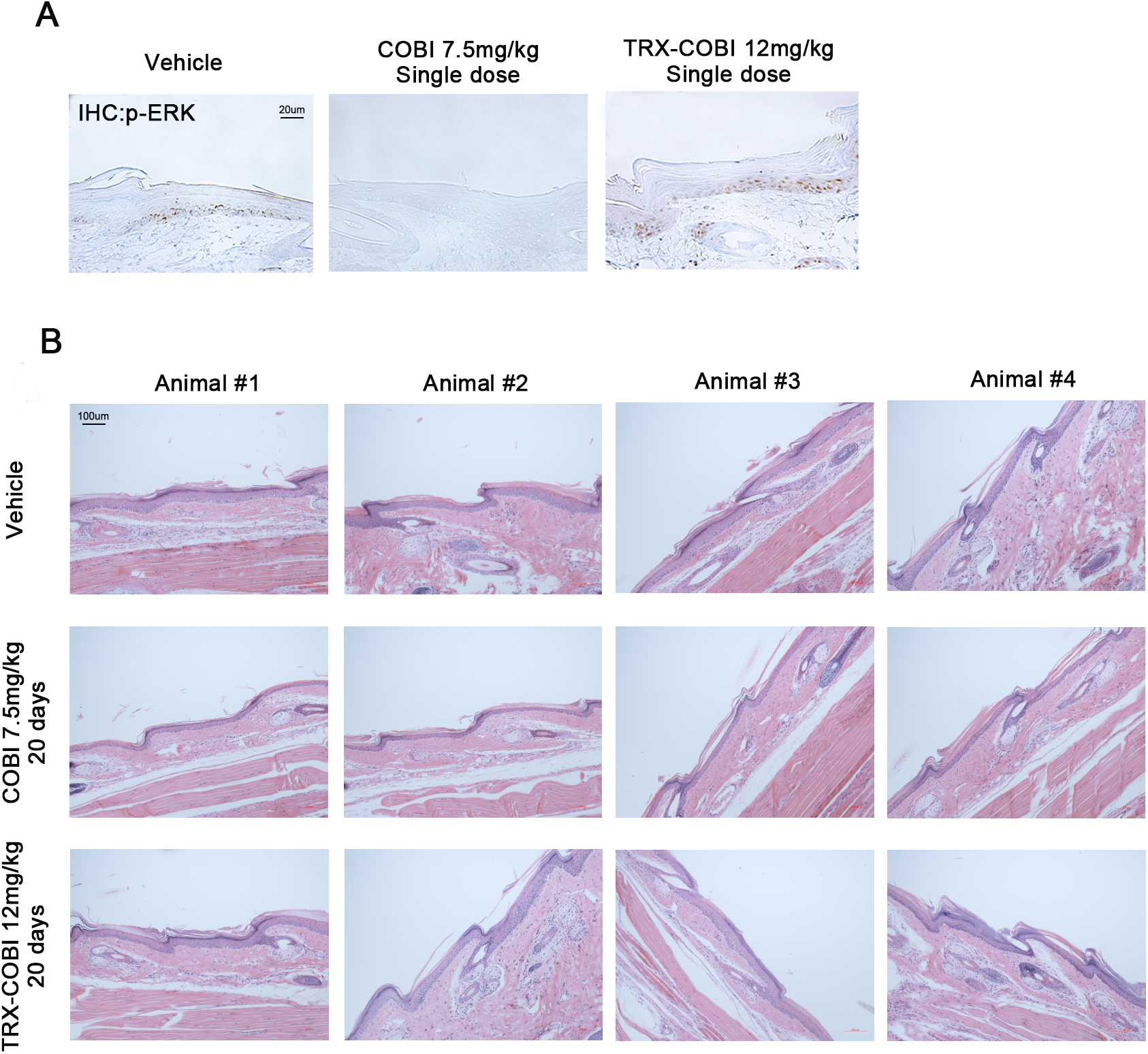
(A) IHC staining with phosphorylated ERK of tail skin (10X) after the single dose of vehicle, the equimolar dose of COBI and TRX-COBI for 8h, corresponding to Figure 5C. (B) Images of HE (10X) show the epidermal layer of mouse tail skin after the treatment with vehicle, the equimolar dose of COBI and TRX-COBI for 20days, corresponding to Figure 5D.

**Supplementary Figure S4.**
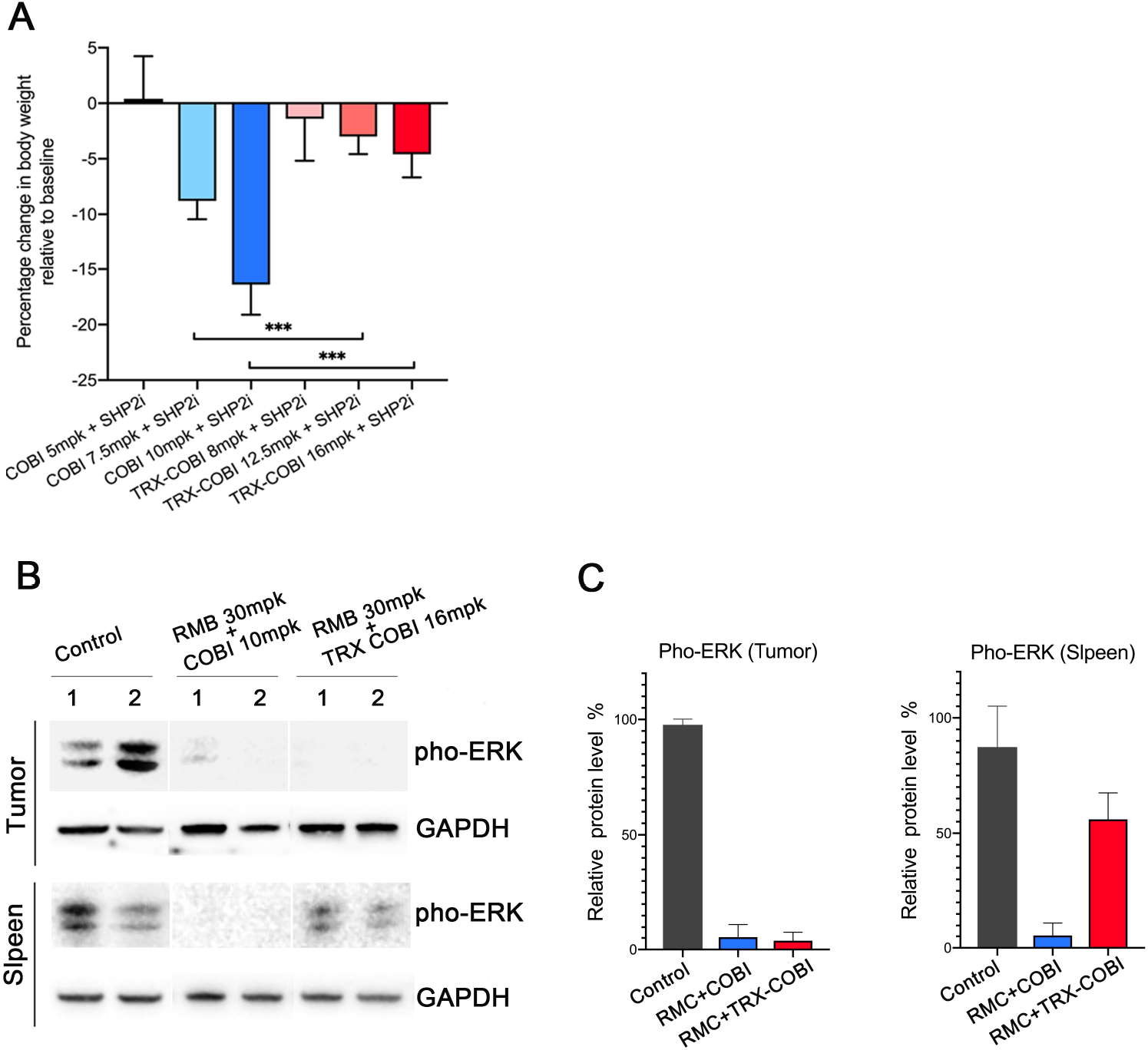
(A) Mouse body weight changes normalized to the baseline after the treatment by the addition of indicated doses of COBI or TRX-COBI to SHP009. Error bars represent mean ± SEM, n=6mice/group and analyzed by two-sample t-test. ***p < 0.001. (B) Western blot for phosphorylated ERK of tumor xenografts and spleen from NSG mice after treatment with vehicle, RMC 30mpk+COBI 10mpk and RMC 30mpk+TRX-COBI 16mpk in the cohort of Figure 6D-E. (C) Quantification of phosphorylated ERK expression in Figure S4B, error bars represent mean ± SEM.

## Acknowledgments

We thank Bob Nichols, Mallika Singh and Trever Bivona for their scientific advice and comments. We also thank PTC and LCA cores at UCSF Helen Diller Comprehensive Cancer Center. This work was supported by NIH, NCI Grants R01 [CA178015, CA222862, CA227807, CA239604, CA230263], U24 [CA210974], U54 [CA224081] grants (EAC). NIH P30CA082103 (ABO). NIH AI105106 and CDRP [W81XWH1810763 and W81XWH1810754] grants (ARR). American Cancer Society Research Scholar Grant (130635-RSG-17-005-01-CCE) (MJE) and the Shorenstein, Rhombauer and Preston Families. Content does not reflect the views of the funders.

## Authors Contributions

Conception and design: H.J, E.A.C, A.R.R.

Development of methodology: H.J, R.K.M, E.A.C, A.R.R.

Data Acquisition: H.J, R.K.M, R.L.G, J.E.K, E.A.C, A.R.R.

Analysis and interpretation of data: H.J, R.K.M, R.L.G, A.B.O, I.Y, B.C.H, N.Z, Y.W, S.C.B, M.J.E, E.A.C, A.R.R

Writing and review of the manuscript: H.J, E.A.C, A.R.R

Study supervision: E.A.C, A.R.R.

## METHODS

### KEY RESOURCES TABLE

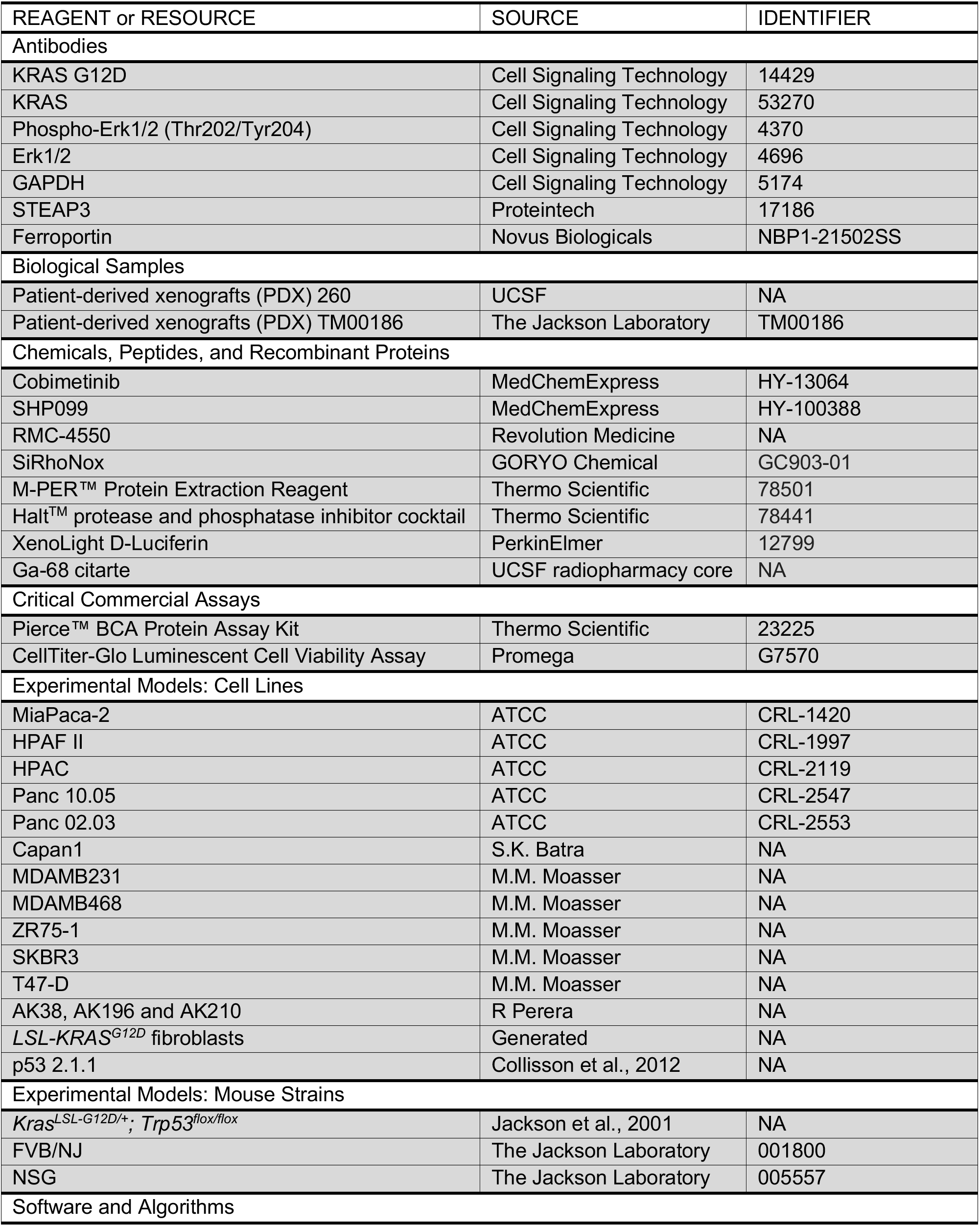
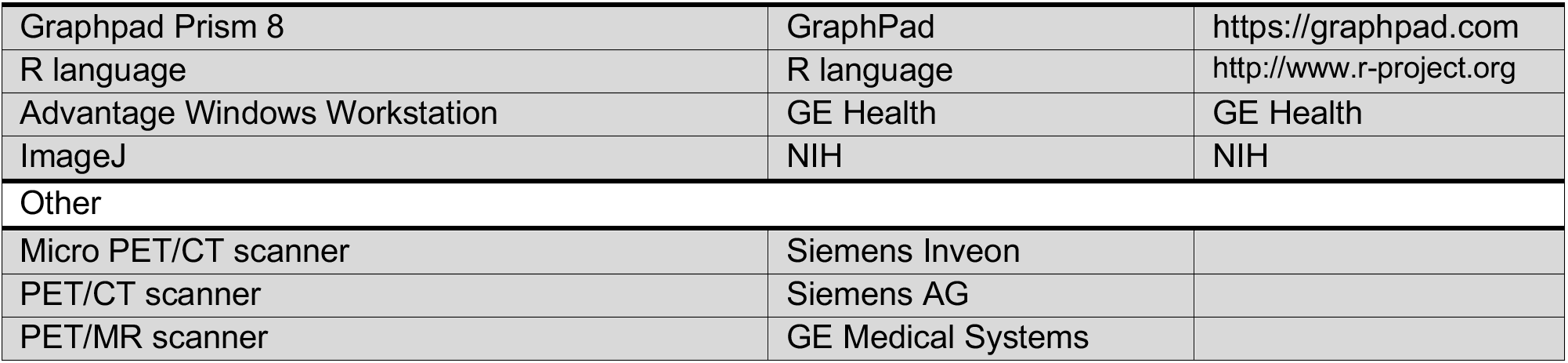

### EXPERIMENTAL MODEL AND SUBJECT DETAILS

#### Cell lines

MiaPaca-2, HPAF II, HPAC, Panc 10.05 and Panc 02.03 are from American Type Culture Collection. Capan1 was provided by S.K. Batra (University of Nebraska, Lincoln, NE). MDAMB231, MDAMB468, ZR75-1, SKBR3 and T47-D were provided by M.M. Moasser (UCSF, CA). *iKras* lines AK38, AK196 and AK210 were provided by R Perera (UCSF, CA). Skin fibroblasts isolation from *LSL-KRAS^G12D^* mouse was performed as described (1). Cells were maintained at 37°C in a humidified incubator at 5% CO2. Cells were grown in appropriate media as recommended by ATCC, and supplemented with 10% fetal bovine serum (Gibco) and 1% penicillin/streptomycin (Gibco). All cell lines tested were negative for mycoplasma contamination.

#### Patient characteristics and accrual

This study was approved by the UCSF Institutional Review Board (IRB) and was compliant with the Health Insurance Portability and Accountability Act. Informed consent was obtained from all patients enrolled into the study.

Patient inclusion criteria were (1) histologically confirmed/clinically suspected pancreatic cancer, (2) At least one lesion ≥ 1cm, (3) Age >18, (4) a signed informed consent indicating that they are aware of the investigational nature of this study, (5) negative for HIV, and (6) patients must not be pregnant or breast feeding.

Patient demographic information including age and sex were recorded as well as their tumor locations and histological grade.

#### Mice models

##### Small animal PET/CT and biodistribution studies

6-week-old male NSG mice were purchased from Jackson lab.

##### Orthotopic pancreas xenografts

Derivation of the p53 2.1.1 FVB/n mice has been described (3).

##### *KPT* lung tumor model

*Kras ^LSL-G12D/+^;p53^flox/flox^;R26^LSL-tdTomato^ (KPT)* mice have been previously described. Mice were bred on a mixed background.

##### PDXs

For generation of the PDX 260 model, informed consent was obtained from the patient as per an open IRB-approved protocol at the University of California, San Francisco. Tumors were implanted in NSG mice (Jackson Lab)

### METHOD DETAILS

#### Reagents and antibodies

The following antibodies were used: KRAS G12D (Cell Signaling Technology, 14429, Dilution: 1:1000); KRAS (Cell Signaling Technology, 53270, Dilution: 1:1000); Phospho-Erk1/2 (Thr202/Tyr204) (Cell Signaling Technology, 4370, Dilution: 1:1000); Erk1/2 (Cell Signaling Technology, 4696, Dilution: 1:1000); GAPDH (Cell Signaling Technology, 5174, Dilution: 1:1000); STEAP3 (Proteintech, 17186, Dilution: 1:1000); Ferroportin (Novus Biologicals, NBP1-21502SS, Dilution: 1:1000). Cobimetinib and SHP099 were purchased from MedChemExpress. RMC-4550 material used in these studies was provided by Revolution Medicines Inc., Redwood City, CA, USA in collaboration with Sanofi, Paris, France.

#### Image protocol

Patients were injected with up to 15 mCi (555 MBq) (average 7.42 mCi [274.6 MBq], range 3.7 to 11.9 mCi [136.9 to 438.5 MBq]) ^68^Ga-citrate intravenously. PET acquisition was acquired between 120 and 263 minutes after injection (average 210 minutes). Images were acquired on either a PET/CT or PET/MR. PET/CT examinations were performed on either a Biograph 16 (Hi-Rez) PET/CT scanner (Siemens AG, Erlangen, Germany) with an integrated PET and 16-MDCT scanner or a Discovery VCT PET/CT scanner (GE Medical Systems, Milwaukee, WI) with an integrated PET and 64-MDCT scanner. A low-dose CT was acquired for PET attenuation correction. PET/MR images were performed on a SIGNA PET/MR (GE Medical Systems, Milwaukee, WI). Attenuation correction for PET reconstruction was performed using a MR-based attenuation correction (MRAC) technique provided by the scanner manufacturer.

#### Image analysis

Maximum intensity projection (MIP), axial, coronal and sagittal reconstructions and PET/MR fused images were reviewed on an Advantage Windows Workstation (AW, Waukesha, WI). PET images were evaluated by trained nuclear medicine physician blinded to the results of conventional imaging scans as well as clinical/genomic features of the case and scored for the presence of PET avid lesions. Lesions were considered PET positive if uptake was focal, greater than the adjacent background soft tissue and not in an expected physiologic structure such as the urinary bladder, vessels or salivary glands.

For semi-quantitative analysis, a volume of interest (VOI) was manually drawn around PET-avid lesions and SUV_max_ were recorded. The location of abnormal radiotracer uptake was compared to CT and nuclear medicine bone-scans. Additionally, SUV_mean_ values were recorded in the liver, paraspinous soft tissues, bone (right sacrum), and mediastinal blood pool for determination of normal structures.

With conventional imaging, soft tissue metastases were considered positive if greater than 1 cm in long axis, except for lymph nodes that were considered positive if greater than 1.5 cm in short axis. Bone lesions on radionuclide scan were considered positive if uptake was focal and not in a pattern consistent with arthritis or antecedent trauma/fracture.

#### Animal studies

All experiments were conducted in the AAALAC accredited University of California, San Francisco in accordance with all applicable local requirements, including approval by the IACUC.

##### Small animal PET/CT and biodistribution studies

6-week-old male NSG mice were purchased from Jackson lab. Mice were inoculated with 10^7^ cells subcutaneously into one flank in a 1:1 mixture (v/v) of media and Matrigel. Tumors were palpable within 20 days after transplantation. 68Ga-citrate was prepared by the Radiopharmacy core facility at UCSF as previously reported (2). 18F-TRX was prepared using the previously developed protocol. Radiotracers were administered via tail vein injection (150-250 µCi/mouse). Imaging data were acquired 4 hours post injection of 68Ga-citrate, and 90 min post injection of ^18^F-TRX. The image acquisition was performed on the Siemens Inveon micro PET/CT under anesthesia by using 2.5% isoflurane at 4 hours post injection. The resulting imaging data was directly reconstructed, decay-corrected, and analyzed by AMIDE software, which was also used to place the region of interest (ROI) to calculate SUV data from the static acquisition. After the imaging data were collected, animals were euthanized by cervical dislocation. Tissues were removed, weighed and counted on a Hidex automatic gamma counter for accumulation activity. The mass of the injected radiotracer was measured and used to determine the total number of CPM by comparison with a standard of known activity. The data were background- and decay-corrected and expressed as the percentage of the injected dose/weight of the biospecimen in grams (%ID/g).

##### Orthotopic pancreas xenografts

Derivation of the p53 2.1.1 FVB/n mice has been described (3). These cells were labelled in vitro with a lentiviral vector encoding a firefly luciferase to give rise p53 2.1.1syn_Luc. 1000 cells were orthotopically implanted in 6-week-old FVB/n mice in 20 μL composed 1:1 mixture (v/v) of media and Matrigel. Mice were treated with 7.5 mg/kg of COBI, 12mg/kg TRX-COBI or vehicle control (saline) by daily IP for 3 weeks. Bioluminescent imaging was performed twice a week to monitor tumor growth.

##### *KPT* lung tumor model

*Kras ^LSL-G12D/+^;p53^flox/flox^;R26^LSL-tdTomato^ (KPT)* mice have been previously described. Mice were bred on a mixed background. Adenovirus expressing Cre recombinase (Viraquest, University of Iowa) were administrated intratracheally as previously described (4). Briefly, 2-3 months old mice were infected with 2X10^7^ or 10^8^ pfu of adenovirus expressing Cre recombinase. After 40 days, 10mg/kg of COBI, 16mg/kg of TRX-COBI or vehicle control (saline) were dosed by IP five days a week for 6 weeks. Mice OS was analyzed and images of tumors expressing tdTomato were acquired with Zeiss Zeiss AxioImager microscope.

##### PDXs

For generation of the PDX 260 model, informed consent was obtained from the patient as per an open IRB-approved protocol at the University of California, San Francisco. Tumors were implanted in NSG mice (Jackson Lab). TM00186 were obtained from Jackson laboratory. Both tumors were used at low passage for the experiments described herein. In figure 4, once tumors reached an average volume of ~300 mm^3^, mice were treated with 7.5 mg/kg of COBI, 12mg/kg TRX-COBI or vehicle control (saline) by daily IP for 25 days. In Figure 6, when tumors reached an average volume of ~300 mm^3^, animals were randomized into six groups as indicated. RMC-4550 was dosed by oral at 30 mg/kg. Measurement of tumor volume and body weight were performed twice a week.

##### TRX-COBI synthesis

The (*R,R*)- and (*S,S*)-TRX-COBI conjugates were prepared by coupling to known TRX intermediates via activated nitrophenyl carbonate as we have previously described (5) and as further detailed in the Supplemental Information.

##### TRX-PURO assay

Automated cell imaging was conducted using a GE IN Cell2000 automated cell imager. Graphing and analysis of data was done in GraphPad Prism 6. 3000 cells per well were plated in 96-well black μClear tissue culture plates (Greiner). The following day, cells were incubated with a mixture of growth medium and adeno-cre viral supernatant for 6 hours. Cells were then exposed to puromycin or TRX-PURO at 1 μM (diluted in cell culture medium from 1,000× DMSO stocks) in medium for 24, 48 and 73 hours before medium was removed and cells were washed with PBS, fixed in 4% PFA for 10 min at RT then washed twice with PBS and once with PBS containing 0.1% Triton X-100. Cells were then stained with Kerafast anti-puromycin antibody (3RH11, 1:500) in PBS with 10% FBS and 0.1% Triton X-100 for 30 min at 37 °C. Cells were washed once with PBS and once with PBS containing 0.1% Triton X-100 then stained with anti-mouse secondary FITC (488 nm excitation, 535 nm emission) antibody (1:100) and Hoechst nuclear stain at a final concentration of 10 μg/mL in PBS with 10% FBS and 0.1% Triton X-100 for 30 min at 37 °C. Cells were washed once with PBS containing 0.1% Triton X-100 and once with PBS then stored in PBS and imaged using an IN Cell 2000 automated cell imager at 10× magnification with 6–9 images per well in FITC and DAPI channel fluorescence. Images were analyzed for nuclei count and puromycin incorporation by IN Cell developer software. Puromycin incorporation was assessed by mean cellular fluorescence density in the FITC channel for each cell as defined by targets seeded with nuclei in the DAPI channel. Average cellular fluorescence density under each condition was determined, and reported values represent the mean average per well across triplicates ± s.e.m. Signal in cells treated with the TRX-PURO was normalized to that in cells treated with free puromycin. These values were then normalized to untreated cells and reported at percent change from untreated.

#### Cell Viability Assays

3000-5000 cells optimized for each cell line were seeded on day 1, drug was added on day 2 (using DMSO normalized to 0.1%), and the cell viability was determined using CellTiter-glo (Promega) on day 4. Viability curves were generated using GraphPad Prism 6.

#### Histological Analysis

HE staining and immunohistochemistry were performed on 4-μm-thick sections of 4% paraformaldehyde-fixed and paraffin-embedded tissues. Tail sections were decalcified by incubation of trimmed paraffin blocks for 10–15 min on paper towels soaked in 1 N HCl. Epidermis thickness was measured by Zeiss AxioImager microscope. Tumors sections were stained with phospho-p44/42 (Thr202/Tyr204, 9101, 1:50).

#### PRISM screening

Cancer cell line profiling was performed using the PRISM platform, as previously described(Yu et al., 2016). Cell treatment and data analysis were performed as described in https://depmap.org/ repurposing. Raw Luminex signal was converted to EC_50_ values for each cell line.

#### Detection of Labile Ferrous Iron

Cells were seeded and cultured overnight. Culture medium was removed on the following day rinse twice gently with PBS. A final concentration of 5 uM of SiRhoNox (FerroFarRed, GORYO Chemical) in a serum-free culture medium was added to the dish and incubate for 1 hour at 37°C. Fluorescence images were then acquired with Zeiss Spinning Disk Confocal microscope. Immunofluorescence was quantified using ImageJ software.

#### Pharmacokinetic Assays

NSG mice were administered a single IP dose of COBI (7.5mg/kg, n=3) or TRX-COBI (12mg/kg, n=3). Blood, tumor, liver and spleen were collected after 7 hours analyzed by Integrated Analytical Solutions, Inc (Berkeley, CA).

#### Western Blotting

Tumor tissue and organ tissues was flash frozen in liquid nitrogen and homogenized in M-PER lysis buffer (Thermo Scientific) plus Halt^™^ protease and phosphatase inhibitor cocktail (Thermo Scientific). Protein concentrations were determined with the Pierce BCA Protein Assay Kit (Thermo Scientific), and extracts were loaded onto NuPAGE Bis–Tris SDS gels and immunoblots were visualized by LiCOR Odyssey system.

#### *STEAP3* knockdown

For stable and lentivirally transfected shRNA-based knockdown experiments, viruses were generated in HEK293T cells transfected with lentiviral packaging vectors along with vectors expressing pGIPZ-shSTEAP3 using Fugene6 (Promega). Two distinct hairpins were chosen for the experiments. Their sequences are as follows: STEAP3-1, 5′-TGAAGAACTTGTTCTGGCT; STEAP3-2, 5′-TGACCACTGTGCAAGTGGG-3′. Viral supernatant collected from confluent monoculture was filtered and used to infect MiaPaca-2 and Capan1 pancreatic cancer cells. A total of 0.5 × 10^6^ cells was seeded in one well of a 6-well chamber and allowed to grow for overnight. The following day, cells were incubated with a 1:2 mixture of growth medium and viral supernatant collected from HEK293T cells. Polybrene was added at 8 µg ml/l.

### QUANTIFICATION AND STATISTICAL ANALYSIS

Statistical tests were performed using GraphPad Prism 7.0 or the R language. Two-sided two-sample t-tests were used for comparisons of the means of data between two groups. One-way ANOVA was used for comparisons among multiple independent groups. The Cox proportional hazards model was used to asses OS, while the Kaplan-Meier method was used to visualize STEAP3 expression when stratified into high versus low. For animal studies, animals were randomized before treatments, and all animals treated were included for the analyses.

